# A sensitive bioassay for detecting *Plasmodiophora brassicae* in canola field soils

**DOI:** 10.64898/2026.07.21.739946

**Authors:** Will Feindel, Kher Zahr, Ronald Nyandoro, Shiming Xue, Tiesen Cao, David Feindel, Michael W. Harding, Habibur Rahman, Fengqun Yu, Jie Feng

## Abstract

We describe a biodegradable-cup bioassay for detecting viable *Plasmodiophora brassicae* in soil samples. Soil samples either artificially inoculated with *P. brassicae* resting spores or collected from canola fields were aliquoted into biodegradable cups containing 20 g of soil per cup. Two cups representing the same soil sample or inoculum concentration were placed in each pot filled with Sunshine Mix. Six seeds of the canola cultivar ‘Westar’ were sown into each cup and thinned to four seedlings per cup ten days after planting. After four weeks, roots were examined for the presence of clubroot galls. Across three independent inoculated-soil experiments, galls were observed in samples containing as few as 1 resting spore g^-1^ soil. In contrast, under a qPCR assay evaluated in parallel, consistent amplification across three technical replicates was obtained only at 100 resting spores g^-1^ soil or greater. In field samples, the bioassay produced galls from 11 qPCR-positive samples and seven of ten qPCR-negative samples. Although the bioassay is not intended for rapid diagnosis or direct quantification, it provides a practical tool for annual clubroot surveys and for studies requiring recovery, propagation, or characterization of viable *P. brassicae* from soil samples collected across diverse geographic regions.

## Introduction

Clubroot, a destructive disease of cruciferous crops caused by the soilborne protist *Plasmodiophora brassicae* Woronin, remains one of the most significant challenges to canola (*Brassica napus*) production in Canada (Feng et al. 2014). The disease was first identified in canola field in Canada in 2003, when infestations were reported in a small number of fields near Edmonton, Alberta (Tewari et al. 2005). Since then, it has spread to more than 4,000 fields in 47 counties and municipal districts in Alberta (Strelkov et al. 2025) and has been confirmed in canola fields in Saskatchewan (Ziesman et al. 2019), Manitoba (Froese et al. 2019), Ontario (Al-Daoud et al. 2018), and North Dakota (Chittem et al. 2014).

The persistence and spread of clubroot are largely attributable to the production of abundant resting spores by *P. brassicae*. These spores can remain viable in soil for as long as two decades (Wallenhammar 1996), serving as the principal inoculum reservoir in infested fields. As a result, estimating resting spore populations in soil has become a key component of clubroot diagnosis and risk assessment. A range of detection techniques has been developed for this purpose, with PCR-based approaches, including conventional PCR, quantitative PCR (qPCR), and digital PCR, being regarded as highly sensitive and reliable tools for spore detection and quantification (Faggian and Strelkov 2009; Wen et al. 2020). In contrast, bioassays, which involve growing susceptible host plants in soil samples and evaluating disease development, have received comparatively little attention as a diagnostic method.

Despite their limited use, bioassays offer several practical advantages. Relative to qPCR, they are generally less expensive and require less specialized laboratory equipment. In addition, bioassays can evaluate a substantially larger soil aliquot than is typically represented in a single DNA extraction and qPCR reaction. By comparison, qPCR analyses are often conducted on DNA extracted from a small subsample of soil, commonly no more than 100 mg, in accordance with the recommendations of many commercial soil DNA extraction kits. The ability to assess a larger proportion of the submitted sample may improve detection when resting spores are unevenly distributed or present near the detection threshold. The fact that bioassays have not been widely adopted for routine clubroot diagnostics is primarily due to three commonly perceived limitations: they require greenhouse facilities and longer testing periods, they are generally unable to provide direct quantitative estimates of resting spore density, and they are often assumed to be less sensitive or less reliable than molecular methods.

Since 2016, the Alberta Plant Health Lab has routinely tested soil samples collected from canola fields across Alberta for the presence and abundance of *P. brassicae* resting spores (Fu et al., 2020). Although probe-based qPCR has been the standard testing approach, characteristics of the submitted samples and the intended use of the results suggested that a bioassay-based method could offer important advantages. Most samples consisted of relatively large soil volumes (100-500 g) collected through annual provincial disease surveys and research programs focused on resistance breeding, population dynamics of the pathogen, and clubroot management. Recovery of viable *P. brassicae* from these samples has been also a demand for downstream applications such as propagation, isolate purification, virulence monitoring, and race characterization using canola differential or single-gene lines (Cao et al. 2026; Zhang et al. 2026).

Furthermore, rapid turnaround times and direct quantitative data were generally not required for these samples. Discussions with diagnostic laboratories in other clubroot- affected provinces indicated that similar types of samples are commonly submitted across Canada (J. Feng, unpublished). Accordingly, the present study was undertaken to evaluate the suitability of a biodegradable-cup bioassay for detecting viable *P. brassicae* in soil samples, with particular emphasis on operational sensitivity, repeatability, and performance with field-collected soils.

## Materials and Methods

### Chemicals and standard procedures

Unless otherwise specified, all chemicals and laboratory instruments were purchased from Fisher Scientific Canada (Ottawa, ON). Primers and probes were synthesized by Integrated DNA Technologies (Coralville, IA). DNA was extracted using the DNeasy PowerSoil Pro Kit (Qiagen Canada, Toronto, ON) on a QIAcube Connect instrument (Qiagen Canada). DNA was eluted in 50 µL of nuclease-free water. When required, DNA concentration was determined using a NanoDrop 1000 spectrophotometer.

### Preparation of Plasmodiophora brassicae inoculum

Clubroot galls from canola cultivar ‘Westar’ were collected from a field nursery (53.6467, −113.3758) at the Alberta Plant Health Lab and stored at −20°C until use. For each inoculation assay, a resting spore suspension was prepared from frozen galls following the method described by Fei et al. (2016). The suspension was serially diluted to a final concentration of approximately 2.5-5 × 10^6^ spores mL^-1^, corresponding to 10-20 spores per 0.04-mm^2^ haemocytometer square, which we believed to be the range that can provide most accurate counting. The final spore concentration was then determined as the mean of independent counts performed by three laboratory members and repeated until the standard deviation was less than 10% of the mean. A series of 10-fold serial dilutions was then prepared, ranging from 1 × 10^7^ to 1 × 10^1^ spores mL^-1^.

### Preparation of *P. brassicae*-inoculated soil samples

A black chernozemic soil (Pennock et al. 2011) was collected from a research field (53.6387, −113.3683) at the Alberta Plant Health Lab. The soil was autoclaved twice at 121°C and 15 psi for 1 h, followed by drying at 50°C for two weeks. The dried soil was homogenized using a laboratory blender and aliquoted into 1.5-mL microcentrifuge tubes (100 mg per tube) and 50-mL centrifuge tubes (20 g per tube). Ten microlitres and 2 mL of each resting spore dilution were added to the 1.5-mL and 50-mL tubes, respectively. The tubes were vortexed thoroughly to ensure uniform distribution of the inoculum throughout the soil. Soil samples in the 1.5-mL tubes were used for DNA extraction, whereas those in the 50-mL tubes were used for the bioassay.

### Preparation of field soil samples

Soil samples were collected from 21 canola fields located within the City of Edmonton and ten counties across Alberta. At each field, soil was sampled using a W-shaped five-point sampling pattern, and the five subsamples were combined into a single composite sample. Two control samples were included in subsequent analyses: (i) an artificially inoculated soil sample containing 1 × 10^5^ spores g^-1^ soil and (ii) a soil sample collected from a clubroot-infested field in Leduc County in 2008 that had been stored in a sealed container at room temperature. All soil samples were air-dried for 48 h, homogenized using a laboratory blender, and aliquoted into 1.5-mL microcentrifuge tubes (100 mg per tube) and 50-mL centrifuge tubes (20 g per tube). Soil samples in the 1.5-mL tubes were used for DNA extraction, whereas those in the 50-mL tubes were used for the bioassay.

### Quantitative PCR assay

Probe-based quantitative PCR (qPCR) was performed using the primers and probe developed by Wallenhammar et al. (2012). Each 20-µL reaction contained 10 µL of 2× PrimeTime Gene Expression Master Mix (Integrated DNA Technologies), 2 µL of DNA template regardless of DNA concentration, 0.25 µM of each primer, and 0.15 µM of probe. The qPCR program consisted of an initial denaturation at 95°C for 2 min, followed by 40 cycles of 95°C for 5 s and 60°C for 30 s. All reactions were performed with technical replicates in triplicate. A sample was recorded as having no signal when none of the three technical replicates generated a quantification cycle (Cq) value.

### Bioassay

Biodegradable seedling cups (4 × 4 × 5 cm) were purchased from Amazon (https://a.co/d/07jENx3n; Fig. S1). Each 20-g inoculated or field soil sample from a 50-mL centrifuge tube was transferred into a biodegradable cup. For each soil sample or inoculum concentration, two cups were prepared and placed in the same 12 × 12 × 12 cm pot filled with Sunshine Mix #4 (Sun Gro Horticulture, Agawam, MA), such that the soil level in the cups was flush with the surrounding Sunshine Mix (Fig. 1a). Six seeds of canola cultivar ‘Westar’ were sown into each biodegradable cup. The pots were kept in a greenhouse maintained at 25°C/22°C (day/night) with a 16-h photoperiod. Ten days after planting, the seedlings were thinned to four plants per cup, resulting in up to eight evaluated plants per soil sample (Fig. 1b). Four weeks after planting, the roots were harvested, washed, and examined for the presence of clubroot galls.

**Fig. 1.**
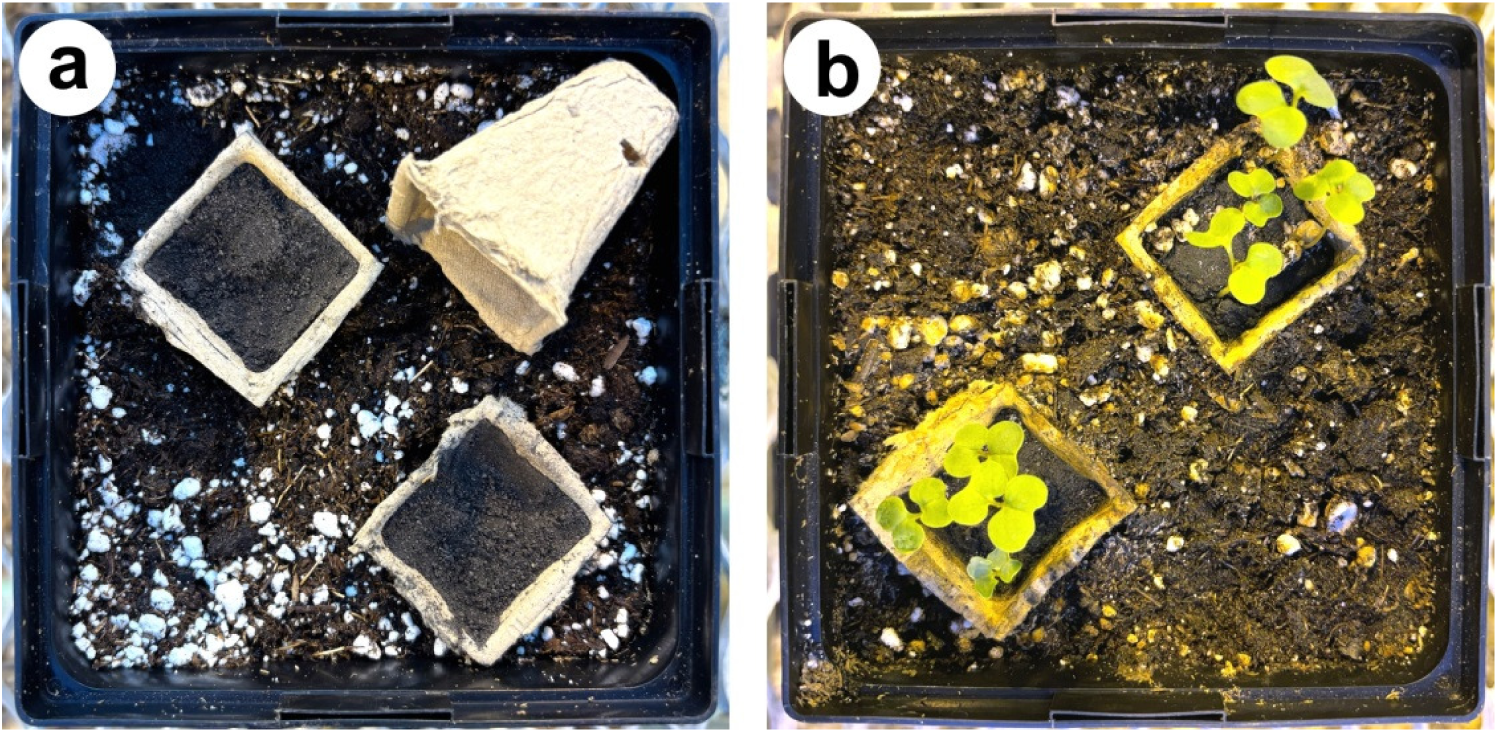
Bioassay setup. a, Twenty grams of soil sample were placed in each biodegradable cup, and two cups were positioned in a plastic pot filled with Sunshine Mix such that the soil level in the cups was flush with the surrounding Sunshine Mix. b, Six canola seeds were sown in each cup, and four seedlings were retained after emergence.

### Evaluation of inoculated and field soil samples

The experiment on the inoculated soil was conducted three times using three sets of inoculated soil samples prepared from separate soil batches and independently prepared resting spore suspensions. In each repeated experiment, a non-inoculated soil treatment was included as a negative control. DNAs extracted from each set of 100-mg soil samples containing serial dilutions of *P. brassicae* resting spores (1 × 10^6^ to 1 × 10^0^ spores g^-1^ soil) were used to generate three qPCR standard curves. Corresponding bioassays were conducted independently using the three sets of 20-g soil samples. For each set of soil samples, the bioassay included three pots as three biological replicates. The pots were arranged in the greenhouse following a randomized complete block design (RCBD). For field soil samples, qPCR assays and bioassays were performed on each sample following the same design as for the inoculated soil; however, the experiment for the bioassay was not repeated, except for the 2008 Leduc County soil sample, which was tested three times.

## Results

### qPCR consistently detected *P. brassicae* at 1 × 10^2^ spores g^-1^ soil or greater

qPCR assays were performed using DNA extracted from soil samples inoculated with serial dilutions of *P. brassicae* resting spores. For soil samples containing resting spores at ≥1 × 10^2^ spores g^-1^ soil, Cq values were obtained consistently from all three technical qPCR replicates. A standard curve was constructed using the Cq values from soil samples containing 1 × 10^2^ to 1 × 10^6^ spores g^-1^ soil (Fig. 2), yielding a calculated qPCR efficiency of 0.94. For samples with spore concentrations below 1 × 10^2^ spores g^-1^ soil, amplification was inconsistent or absent across the three technical replicates. Thus we concluded that the detection limit of the qPCR was 1 × 10^2^ spores g^-1^ soil.

**Fig. 2.**
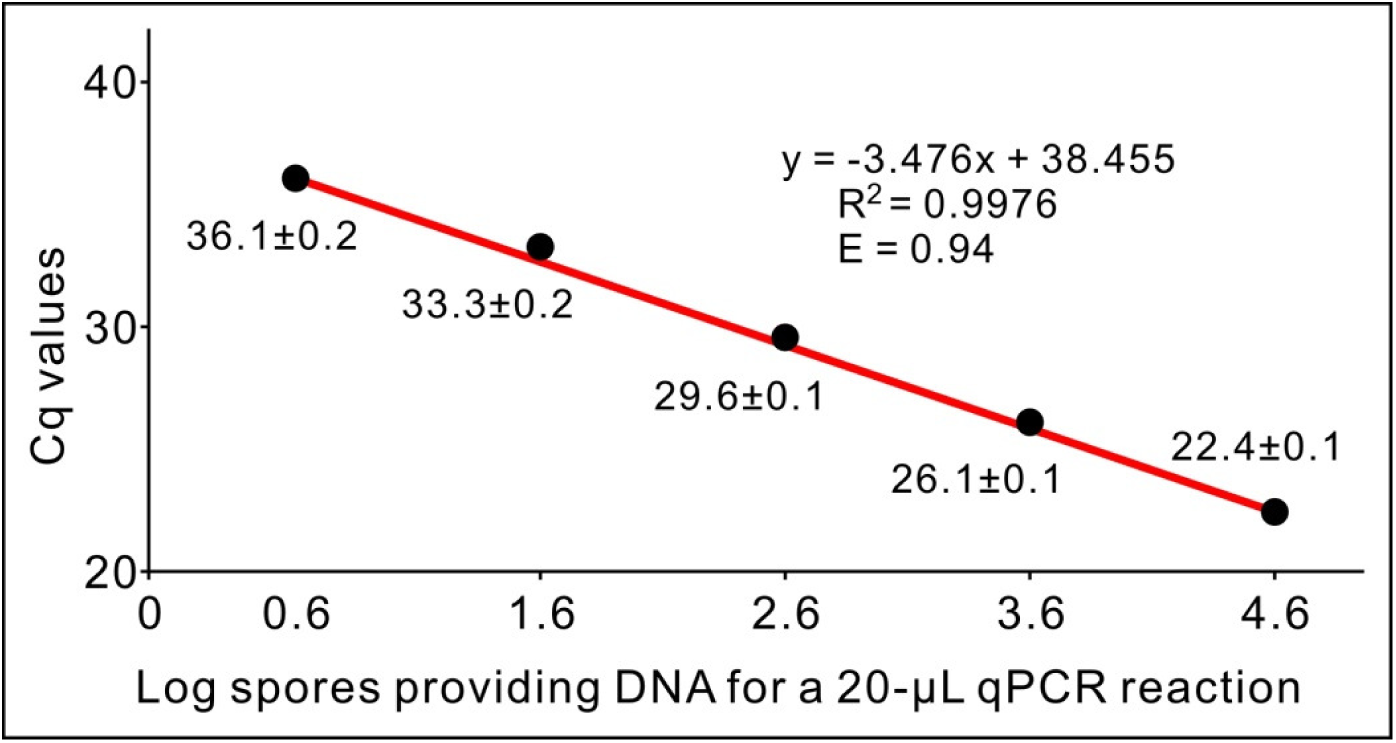
Standard curve of a probe-based qPCR system using DNA extracted from soil samples inoculated with serial dilutions of *Plasmodiophora brassicae* resting spores. The curve was generated from mean quantification cycle (Cq) values plotted against the log_10_ of calculated spore-equivalent DNA input per 20-µL qPCR reaction. The R^2^ value of the equation and primer efficiency (E) are indicated on the curve. Efficiency was calculated as E = −1 + 10^(−1/slope)^. Each data point represents the mean of three technical replicates ± standard deviation.

### The biodegradable-cup bioassay detected viable *P. brassicae* at 1 spore g^-1^ soil

Bioassays were performed using soil samples inoculated with serial dilutions of *P. brassicae* resting spores. Four weeks after planting, all cups were penetrated by canola roots and partially degraded (Fig. 3a). No clubroot galls were observed in the non-inoculated control. Clubroot galls were observed on plants grown in inoculated soil samples, including those containing 1 spore g^-1^ soil (Fig. 3b). For soil samples containing ≥1 × 10^3^ resting spores g^-1^ soil, clubroot galls developed consistently on all evaluated plants. For soil samples containing <1 × 10^3^ resting spores g^-1^ soil, clubroot galls developed on a subset of the evaluated plants. These results demonstrate that the bioassay is capable of detecting *P. brassicae* at concentrations as low as 1 spore g^-1^ soil.

**Fig. 3.**
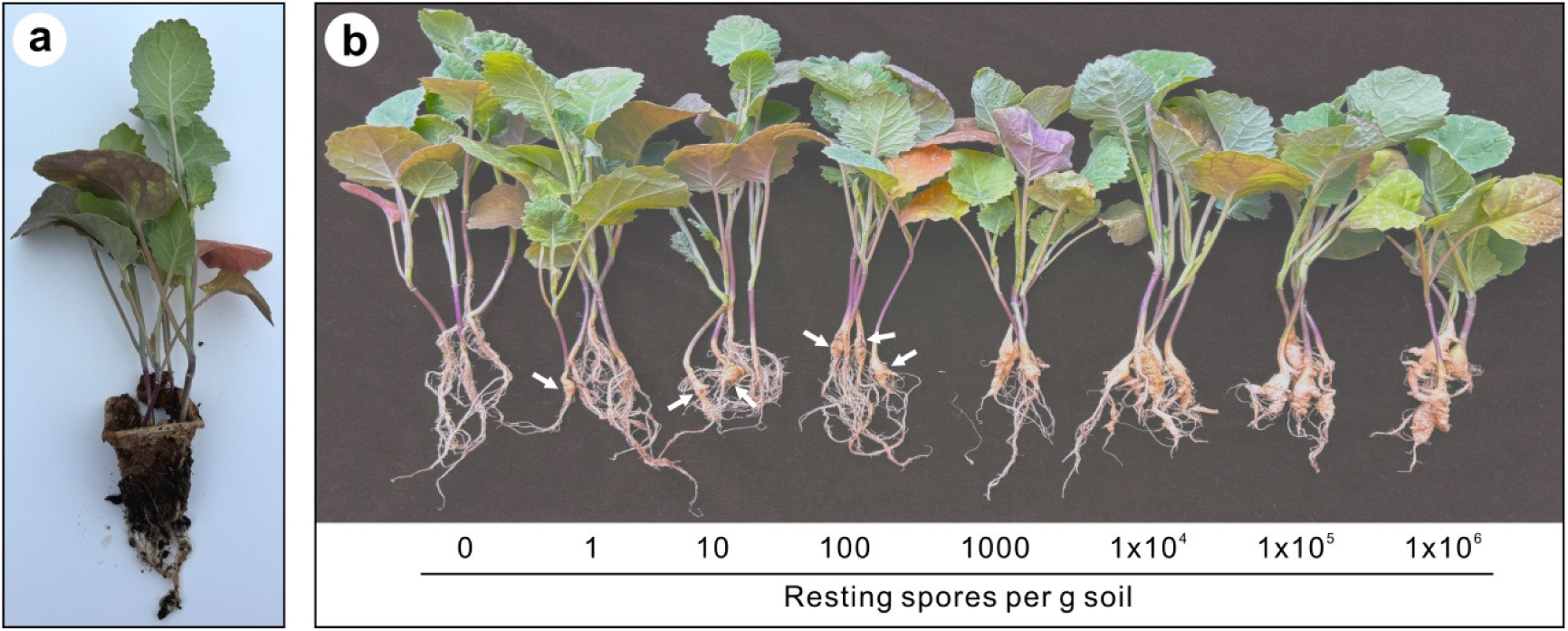
Clubroot development on canola plants four weeks after planting in the bioassay. a, A canola plant grown in non-inoculated soil, showing root penetration through the biodegradable cup. b, Root galls on canola plants grown in soil samples inoculated with serial dilutions of *Plasmodiophora brassicae* resting spores. Plants in one biodegradable cup are shown for each inoculum concentration. Arrows indicate galls when not all plants within a cup exhibited symptoms.

### Field soil samples supported the high operational sensitivity of the bioassay

The qPCR and the bioassay were evaluated using 21 soil samples collected in 2025, one soil sample collected in 2008 (tested in triplicate), and one artificially inoculated soil sample used as a positive control (Table 1). Of the 21 field samples collected in 2025, 11 generated qPCR Cq values, whereas ten generated no signal. All 11 qPCR-positive samples developed clubroot galls in the bioassay. Among the ten samples that did not generate a qPCR signal, seven nevertheless produced clubroot galls in the bioassay.

**Table 1.** qPCR and bioassay results for soil samples collected from canola fields.

| Sample ID | Origin | Cq of qPCR<sup>a</sup> | Galls<sup>b</sup> |
| --- | --- | --- | --- |
| 1 | Athabasca County | 35.5 ± 0.2 | 7 |
| 2 | Camrose County | 19.3 ± 0.1 | 8 |
| 3 | Edmonton | 27.2 ± 0.1 | 8 |
| 4 | Lamont County | No signal | 4 |
| 5 | Leduc County | 19.3 ± 0.1 | 8 |
| 6 | County of Minburn | 28.8 ± 0.1 | 8 |
| 7 | Red Deer County | 33.3 ± 0.1 | 8 |
| 8 | Red Deer County | 31.4 ± 0.2 | 8 |
| 9 | Strathcona County | No signal | 2 |
| 10 | Strathcona County | No signal | 5 |
| 11 | Strathcona County | 38.4 | 3 |
| 12 | Strathcona County | 36.9 ± 0.4 | 6 |
| 13 | Strathcona County | 37.2 ± 0.3 | 2 |
| 14 | Strathcona County | No signal | 0 |
| 15 | Strathcona County | No signal | 0 |
| 16 | Westlock County | No signal | 3 |
| 17 | Westlock County | 31.8 ± 0.1 | 8 |
| 18 | Westlock County | No signal | 0 |
| 19 | Westlock County | No signal | 2 |
| 20 | County of Vermilion River | No signal | 5 |
| 21 | County of Vermilion River | No signal | 3 |
| 22a | Leduc County | 20.5 ± 0.1 | 8 |
| 22b | Leduc County | 21.0 ± 0.1 | 8 |
| 22c | Leduc County | 20.6 ± 0.1 | 8 |
| CK | 1 × 10<sup>5</sup> spores g<sup>-1</sup> soil | 26.3 ± 0.1 | 8 |

<sup>a</sup>DNA was extracted from 100 mg of soil, eluted in 50 µL of water, and 2 µL was used in each qPCR reaction. No signal indicates that no Cq value was obtained from any of the three technical replicates within 40 cycles.
<sup>b</sup>Number of plants with clubroot galls out of eight evaluated plants in the two biodegradable cups.

Both the 2008 soil sample and the inoculated positive control generated Cq values in qPCR and produced clubroot galls in the bioassay. Collectively, these results demonstrate that the bioassay is more sensitive than the qPCR assay for detecting *P. brassicae*, particularly in soil samples with low resting spore concentrations.

## Discussion

The present study demonstrates that a simple biodegradable-cup bioassay can detect viable *P. brassicae* in soil at nominal concentrations as low as 1 resting spore g^-1^ soil. Under the conditions evaluated, the bioassay consistently detected the pathogen at lower inoculum levels than the probe-based qPCR used for comparison, which reliably detected *P. brassicae* only at ≥10^2^ resting spores g^-1^ soil. The field evaluation further supported this difference in operational sensitivity, as clubroot galls developed from several field samples that produced no detectable qPCR signal. Collectively, these results indicate that the biodegradable-cup bioassay provides a sensitive and practical approach for detecting viable *P. brassicae* inoculum in soil and complements existing molecular diagnostic methods.

The higher operational sensitivity of the bioassay is most likely explained by fundamental differences in sampling and detection rather than by analytical performance alone. In the qPCR used in this study, DNA was extracted from only 100 mg of soil, and only a small fraction of the recovered DNA was subsequently analyzed in each PCR reaction.

Consequently, when resting spores are present at very low densities or are unevenly distributed within a soil sample, stochastic effects associated with subsampling, DNA extraction, and PCR amplification may reduce the probability of detection. In contrast, each bioassay evaluated 20 g of soil, increasing the likelihood that viable resting spores were represented in the assay. Furthermore, infection of susceptible canola plants provides an effective biological amplification step, whereby even a small number of viable resting spores can produce readily visible clubroot symptoms. The confined soil volume within the biodegradable cup may also increase opportunities for root-pathogen contact, although this possibility warrants further investigation. Together, these factors provide a biologically plausible explanation for the markedly higher sensitivity observed in the bioassay.

The qPCR detection threshold observed in this study is consistent with previous reports. All PCR- and qPCR-based assays developed for *P. brassicae* target the ribosomal DNA region (Fu et al. 2026), and reported detection limits from soil generally range from approximately 10^2^ to 10^3^ resting spores g^-1^ soil, depending on DNA extraction procedures, assay chemistry, and soil characteristics (Faggian et al. 1999; Cao et al. 2007; Rennie et al. 2011; Wallenhammar et al. 2012; Li et al. 2013; Xing et al. 2021). The consistent detection limit of 10^2^ resting spores g^-1^ soil obtained here therefore agrees well with the published performance of molecular assays and suggests that the superior sensitivity of the bioassay primarily reflects differences in sample representation and biological amplification rather than shortcomings of the qPCR method itself.

An important distinction between qPCR and biological detection methods is that qPCR detects pathogen DNA regardless of viability, whereas the bioassay detects only resting spores capable of infecting host plants. Although this difference means that the two approaches measure different aspects of pathogen presence, it does not diminish the practical value of the bioassay for many applications. First, the detection of viable inoculum is more relevant than the detection of residual pathogen DNA when the objective is to assess disease risk or recover living pathogen populations for further study. Second, despite targeting only viable resting spores, the bioassay demonstrated greater operational sensitivity than the qPCR evaluated in this study, with seven of ten qPCR-negative field samples nevertheless producing clubroot galls. Finally, the successful recovery of viable *P. brassicae* from a soil sample collected in 2008 is consistent with previous reports that resting spores can remain viable in soil for up to 20 years (Wallenhammar 1996), suggesting that most spores in a submitted soil sample are alive and thus the bioassay can be used for detection. Collectively, these observations indicate that biological detection methods provide valuable information that complements molecular diagnostics, particularly when the objective is to determine whether soil contains infectious inoculum rather than simply pathogen DNA.

Although the bioassay requires greenhouse facilities and approximately four weeks to complete, these requirements are unlikely to be limiting for many of the applications for which the assay is intended. Annual clubroot surveys, epidemiological studies, resistance screening programs, and investigations of pathogen population biology generally prioritize detection sensitivity and recovery of viable pathogen material over rapid turnaround time. Because infected plants generated by the bioassay provide living inoculum, the assay also facilitates downstream applications such as pathogen propagation, isolate purification, virulence monitoring, and race characterization using differential or single-gene canola lines (Cao et al. 2026; Zhang et al. 2026). In addition, the assay requires only standard greenhouse facilities and inexpensive materials, making it accessible to laboratories without specialized molecular diagnostic equipment. Several limitations of this study should nevertheless be acknowledged. Field samples were evaluated without biological replication, only one qPCR was included for comparison, and the low-inoculum experiments demonstrate detection of nominal resting spore concentrations rather than a statistically defined limit of detection. Additional studies involving a wider range of soil types, environmental conditions, and molecular detection platforms, together with replicated analyses of detection probability, would provide a more comprehensive assessment of assay performance. Further investigation of the mechanisms contributing to the high sensitivity of the biodegradable-cup system would also be valuable.

In conclusion, the biodegradable-cup bioassay provides a practical and highly sensitive method for detecting viable *P. brassicae* in soil. By evaluating substantially larger soil samples than conventional molecular assays and relying on biological amplification through host infection, the assay achieved greater operational sensitivity than the qPCR workflow evaluated in this study. Although it is not intended to replace molecular diagnostics in situations requiring rapid results or quantitative estimates of resting spore density, it represents a valuable complementary tool for clubroot surveillance, epidemiological investigations, and research requiring the recovery and propagation of viable *P. brassicae* populations.

## Acknowledgements

This study was partially supported by grants from Saskatchewan Agriculture Development Fund and SaskOilseeds to FY.

---

**Fig. S1.**
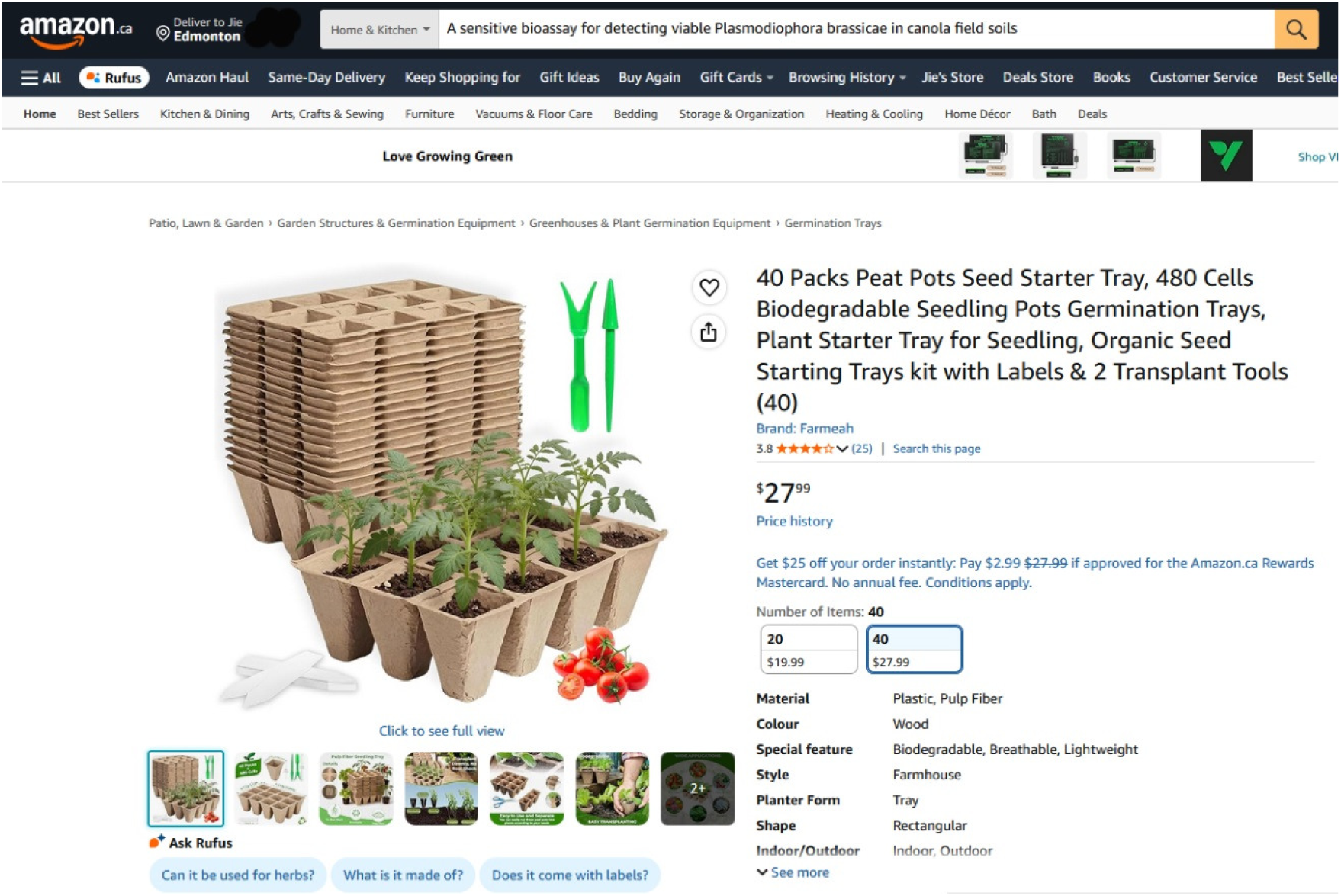
Screenshot of the Amazon product page for the biodegradable seedling cups used in this study. The page was accessed on July 21, 2026.

